# *Legionella* and *Mycobacterium* populations exhibit geographic structuring across and within drinking water systems

**DOI:** 10.64898/2026.01.13.699378

**Authors:** Huanqi He, Samantha DiLoreto, Jinhao Yang, Patrick Milne, Christopher A. Impellitteri, Aron Stubbins, Kelsey Pieper, Katherine Graham, Ching-Hua Huang, Ameet Pinto

**Affiliations:** School of Science and Engineering, Benedict College, Columbia, South Carolina, USA; School of Civil and Environmental Engineering, Georgia Institute of Technology, Atlanta, Georgia, USA; Department of Chemistry and Chemical Biology, Northeastern University, Boston, Massachusetts, USA; The Water Tower, Buford, Georgia, USA; Department of Marine and Environmental Sciences, Northeastern University, Boston, Massachusetts, USA; Department of Civil and Environmental Engineering, Northeastern University, Boston, Massachusetts, USA; Department of Civil and Environmental Engineering, University of North Caroline Charlotte, North Carolina, USA; School of Earth and Atmospheric Sciences, Georgia Institute of Technology, Atlanta, Georgia, USA

**Keywords:** Legionella, Mycobacterium, opportunistic pathogens, disinfection by-products, drinking water distribution systems

## Abstract

Opportunistic pathogens (OPs) within the *Legionella* and *Mycobacterium* can persist and sometimes proliferate in drinking water systems and pose a risk to public health. Most prior research has focused on isolated system components of the drinking water treatment and distribution system and has rarely examined spatiotemporal dynamics across the entire source water, treatment process, and distribution system continuum. This study addresses this critical knowledge gap by quantitative profiling of microbial communities with full length 16S rRNA gene sequencing and flow cytometry, and associated water chemistry parameters, including disinfection byproducts (DBPs), across five full-scale utilities. These utilities reflect varying source water types, geographic locations, treatment regimes, and climate zones. Microbial communities, including *Legionella* and *Mycobacterium* populations, in distribution system were shaped by source water type and exhibited significant community divergence across utilities. Within the same genus, strain-level analyses revealed highly distinct *Legionella* and *Mycobacterium* sequence variants unique to each utility. Interestingly, a substantial proportion of *Legionella* and *Mycobacterium* amplicon sequence variants were both utility specific and often specific to locations within the distribution system, indicating strong geographic structuring both across and within drinking water systems. Understanding the mechanistic underpinnings of this geographic structuring is critical to develop robust strategies for managing and monitoring *Legionella* and *Mycobacterium* populations in drinking water systems.

## 1. Introduction

Opportunistic pathogens (OPs) within the genera *Legionella* and *Mycobacterium* are natural inhabitants in drinking water systems and can persist and proliferate in the drinking water distribution systems (DWDS)^1,2^. Exposure to these pathogens poses serious health risks, particularly for immunocompromised individuals. *Legionella pneumophila* is the primary causative agent of Legionnaires’ disease, a severe form of pneumonia that is fatal for 1 in 10 cases^3^. Clinically relevant species of non-tuberculous mycobacteria (NTM) predominately cause a range of chronic pulmonary infections, with infections due to *Mycobacterium avium* and *Mycobacterium abscessus* being the most common and difficult to treat ^4,5^. Reported infection prevalence from both pathogens has significantly increased over the last two decades^6^, contributing to estimated $2.39 billion annual healthcare costs in the US alone^7^. A wide variety of factors can influence the occurrence of *Legionella* and *Mycobacterium* in drinking water systems^8–10^, including abiotic (e.g., disinfectants, temperature) ^11–14^ and biotic factors (e.g., interactions with protists)^15–17^. Notably, both genera are biofilm formers, and biofilm matrix provides protection from disinfectants and can facilitate long-term colonization in the DWDS^18,19^. Many of the previous studies on *Legionella* and/or *Mycobacterium* have focused on isolated system components where biofilms are prevalent, such as building premise plumbing ^20–22^, distribution mains ^2,23,24^, and water storage tanks ^25,26^. While these studies have yielded critical insights, they often overlook critical upstream segments of drinking water systems, including source water and finished water at the point-of-entry (POE) into the DWDS. These upstream locations establish the microbial and chemical baseline that shapes microbial community assembly.

Understanding how microbial diversity and composition vary across geographic scales is essential to enable robust microbiological monitoring efforts for drinking water systems. The occurrence of certain species is restricted to specific geographic regions (i.e., geographic separation) in aquatic ecosystems reflects the combined influences of environmental selection ^28,29^, dispersal limitation ^29^, and historical contingency across hydrologic systems ^30^. Geographic structuring has been documented in both source water habitats ^31–33^ and engineered water infrastructures ^34–36^. Roeselers et al. observed highly distinct community profiles in samples originating from different DWDS in Netherlands, despite broadly similar treatment processes^36^. Similarly, Ma et al. demonstrated a negative correlation between tap water community similarity and geographical distance, indicating increasing community divergence with spatial separation and that microbial turnover was largely driven by local source water, runoff, and rainfall events^35^. Such studies suggest that the drinking water microbiome, even after strong selective pressures imposed by treatment processes, continue to be shaped by localized environmental conditions. While these and other prior studies have substantially advanced our understanding of the biogeography of drinking water microbiome, there has been little research on fine-scale variations within *Legionella* and *Mycobacterium* populations. This limitation is particularly important given that the strain-level variants within OP genera, i.e., microdiversity, can influence their virulence and persistence in the oligotrophic environment ^37^. As a result, geographic structuring of *Legionella* and *Mycobacterium* populations remains a critical knowledge gap that is directly relevant to microbial risk management.

In this study, we address this critical knowledge gap by characterizing the spatiotemporal trends in five full-scale U.S. drinking water utilities that span variable source waters, treatment configurations, climate conditions, and geographic regions. Our objectives were to: (1) characterize the spatiotemporal variations in drinking water quality and microbial community composition; (2) examine occurrence dynamics of *Legionella* and *Mycobacterium* along the source-treat-distribution continuum; and (3) investigate the geographic structuring of *Legionella* and *Mycobacterium* populations within and across drinking water systems.

## 2. Materials and methods

### 2.1 Description of drinking water utilities and sampling campaign

This study included five full-scale drinking water utilities across different geographic regions of the United States, each with distinct source waters and DWDS sizes. All utilities applied monochloramine at the POE of their DWDS as a secondary disinfectant. Sampling at each utility included three locations: (1) the source water(s), (2) treated water at the POE, and (3) sites within the DWDSs representing a range of water ages. Two sampling campaigns were conducted per utility between January and May 2024, with each site sampled in duplicate. An overview of each utility and the sampling campaign is presented in Table 1, while Table S1 provides details on treatment configurations and their climate zones. For each sampling campaign, 500 mL of source water and 2000 mL of POE and DWDS samples were collected in sterile Nalgene^TM^ PPCO bottles (Thermo Fisher Scientific, MA, USA) containing 10 mg sodium thiosulfate for microbial analyses. An additional 1000 mL was collected at each site in acid-washed and rinsed HDPE bottles for dissolved organic matter analysis (DOM), and 125 mL was collected in HDPE bottles for anion analysis. All samples were shipped overnight on ice to analyzing laboratories for respective analysis.

**Table 1:** An overview of each drinking water utility and the sampling campaign.

| Utility | Source water type | Water production capacity (Million gallons per day – MGD) | Disinfectant | Sampling sites | Sampling dates |
| --- | --- | --- | --- | --- | --- |
| **S1** | Surface | 200 MGD | Primary: O<sub>3</sub>;<br>Secondary: NH<sub>2</sub>Cl | Surface water (n=1),<br>POE water (n=1),<br>DWDS water (n=4) | Winter: Feb-24<br>Spring: May-24 |
| **S2** | Surface | 96 MGD | Primary: HOCl;<br>Secondary: NH<sub>2</sub>Cl | Surface water (n=1),<br>POE water (n=1),<br>DWDS water (n=4) | Winter: Feb-24<br>Spring: May-24 |
| **G1** | Ground | 50 MGD | Primary: HOCl;<br>Secondary: NH<sub>2</sub>Cl | Ground water (n=1),<br>POE water (n=1),<br>DWDS water (n=4) | Winter: Jan-24<br>Spring: Apr-24 |
| **G2** | Ground | 3 MGD | Primary: NH<sub>2</sub>Cl;<br>Secondary: NH<sub>2</sub>Cl | Ground water (n=1),<br>POE water (n=1),<br>DWDS water (n=1) | Winter: Jan-24<br>Spring: Apr-24 |
| **B1** | Surface/gro<br>und<br>variable<br>blend | 14 MGD | Primary: O<sub>3</sub>;<br>Secondary: NH<sub>2</sub>Cl | Blended water (n=1),<br>POE water (n=1),<br>DWDS water (n=4) | Winter: Jan-24<br>Spring: May-24 |

### 2.2 Chemical analyses

Total chlorine residual, water temperature, and pH for all samples were measured and reported by each utility. Ammonia was measured using Hach method 10023 (Hach Company). Dissolved organic carbon (DOC) and total dissolved nitrogen (TDN) were analyzed using a Shimadzu TOC-L+TN analyzer. Anions (i.e., Nitrite, Nitrate, Chloride, and Flouride) were analyzed using EPA Method 300.1 Parts A and B. A suite of DBPs including trihalomethanes (THMs), iodinated THMs (I-THMs), haloketones (HKs), haloacetonitriles (HANs), trichloronitromethane (TCNM), haloacetamides (HAMs), haloacetic acids (HAAs), iodinated HAAs (I-HAAs), and nitrosoamines (NISAMs) were measured as outlined in DiLoreto et al ^39^.

### 2.3 Flow cytometric analyses

Samples were processed in triplicate on Cytoflex Flow Cytometer (Beckman Coulter) to determine total cell counts (TCC) and intact cell counts (ICC). Samples were processed as outlined in Vosloo et al ^22^. Briefly, samples were stained with Invitrogen™ SYBR Green I (SG) (1:100 SG diluted in 10 mM Tris-HCl (pH 8.5, Bioworld) combined with Molecular Probes™ propidium iodide (PI) (3 μM final concentration) at 12 μL SGPI double stain per mL of sample ^22^. Post staining, the samples were incubated in the dark for at least 15 minutes. Two blanks and two negative controls were processed in parallel with the samples. These included (1) unstained DNase/RNase-Free Distilled Water (Thermo Fisher Scientific), (2) stained DNase/RNase-Free Distilled Water, (3) stained Millex-GP filtered sample (0.22 μm pore size, Millipore); (4) stained boiled sample (100°C water bath for 30 minutes). The flow cytometric measurement was performed with a 50mW solid-state laser emitting light at a fixed wavelength of 488 nm. Green and red fluorescence intensities were collected in the B525 channel (525 ± 40 nm) and B690 channel (690 ± 50 nm), respectively, along with sideward (SSC) and forward (FCS) scatter light intensities. All data were processed with the FlowJo software (FlowJo LLC), and electronic gating with the software was used to separate positive signals from noise ^40^.

### 2.4 DNA extractions, full-length 16S rRNA gene sequencing, and data processing

2000 mL POE and DWDS samples and 500 mL of source water samples were concentrated using sterile Sterivex ^TM^ filter units (0.22 μm pore size, Millipore). Genomic DNA was extracted from the filter membranes following a previously described protocol^18^. Negative controls including reagent blank and membrane filter treated with UltraPure™ DNase/RNase-Free Distilled Water (Thermo Fisher Scientific) were processed in parallel for each batch of samples. The library preparation and sequencing were performed by the Functional Genomics Laboratory at the University of Illinois (IL, USA). The full-length 16S rRNA genes were amplified with the KAPA HiFi Hot Start ReadyMix PCR Kit (KAPA Biosystems) and a primer set of 27F (AGRGTTYGATYMTGGCTCAG) and 1492R (RGYTACCTTGTTACGACTT). A total of 12 negative controls and 2 distilled water controls were sequenced in parallel with the samples. Polymerase chain reaction (PCR) conditions were set as follows: denaturation of 95 °C for 3min, 20 cycles at 95 °C for 30s, 57 °C for 30s, and 72 °C for 60s. The 16S amplicons were generated with the barcoded Full-Length Kinnex 16S primers from PacBio and the 2x Roche KAPA HiFi Hot Start Ready Mix. The DNA was converted to a library with the SMRTBell Prep kit 3.0. Sequencing was performed with SPARQ chemistry in a REVIO instrument (Pacific Biosciences) with a 30-hour movie, and circular consensus sequence (CCS) analysis was performed on the instrument. Demultiplexing was done using SMRTlink 25.1 using the following parameters: ccs --min-passes 3 --min-rq 0.999. Raw sequences were processed with DADA2 v. 1.30.0 ^41^ in R v. 4.3.3 ^42^ to trim primers, denoise and merge reads to construct amplicon sequence variants (ASVs) followed by chimera removal using standard workflow. The SILVA nr v.138.1 database ^43^ was used for taxonomic assignment of ASVs within DADA2 and non-bacterial sequences were removed. Contaminating ASV sequences were identified and removed following the Decontam pipeline ^44^. Table S2 summarizes the reads per sample at different stages of data processing in DADA2.

### 2.5 Data analyses

After normalizing samples to equal number of sequence reads, alpha-diversity (Shannon index) and beta-diversity (Bray-Curtis index) analyses were conducted using the phyloseq v. 1.46.0 ^45^ and vegan v. 2.6.6.1^46^ packages. Permutational multivariate analysis of variance (PERMANOVA) was performed using the adonis function in vegan to compare the microbial community composition across different categories. Distance-based redundancy analysis (dbRDA) was used to quantify the variance in the beta-diversity that could be explained by the measured environmental variables using the vegan package. Spearman correlation was used to analyze the relationship between measured variables. ASVs classified to *Mycobacterium* and *Legionella* genera were extracted from unnormalized sequence reads table to calculate their relative abundances. The *Mycobacterium* and *Legionella* ASV sequences were aligned with mafft ^47^, and maximum likelihood method in IQ-TREE v. 2.0.3 ^48^ was used to construct the phylogenetic tree (number of bootstraps = 1000). DESeq2 v. 1.42.1 ^49^ was used on the unnormalized read count table for differential abundance analysis and p-values were adjusted using the Benjamin and Hochberg method^50^. The fast expectation-maximization for microbial source tracking (FEAST) approach^51^ was used to assess the contribution of various sources towards microbial communities in DWDS samples. FEAST employs a Bayesian approach to estimate the proportion of user-defined “source” microbial communities in a given “sink” community. In this study, source water and POE samples were designated as the “sources”, while downstream DWDS samples were considered the “sinks”. For all statistical analyses, the threshold of p < 0.05 was considered significant. Except for IQ-TREE, all other analyses were performed in R v. 4.3.3 ^42^.

## 3. Results and Discussion

### 3.1 Biogeographic structuring and source water characteristics influence water chemistry and microbial community

Water quality measurements for the five utilities are summarized in Table S3. Total chlorine residuals were higher in utilities using ground water as source (3.6±0.4 mg/L) as compared to those blending surface and ground water (2.9±0.4 mg/L) or those using surface water (1.6±0.7 mg/L). Water temperature was elevated for groundwater utilities (26.3±2.4 °C) relative to those with source and ground water blends (15.3±3.4 °C) or surface water utilities (11.2±3.8 °C). This elevated temperature in groundwater systems can be attributed to their warmer climate zones and could also explain the need for a higher chlorine residual, which may be used to counteract the higher rates of chlorine decay at warmer temperatures^8^. Utility G1 consistently had higher DOC (4.6±0.15 mg/L) compared with other utilities (2.3±0.51 mg/L), reflecting the elevated DOC concentration in its source water (7.2±0.61 mg/L). Sulfate and chloride concentrations ranged widely from 0.6 to 58.6 mg/L and 2.6 to 144.5 mg/L across all drinking water samples, respectively. Utility B1 exhibited the highest sulfate (49.4 ± 5.3 mg/L), chloride (118.8 ± 11.0 mg/L), and nitrate (2.4±1.5 mg/L) levels, while the utility S2 had the lowest. These differences largely reflect water chemistry of the source water for each utility.

Variance partitioning analysis revealed that the largest factor contributing to the overall variance in beta-diversity was the location (i.e., different utility IDs), explaining 33.1% of the total variance (ANOVA, p = 0.001). Köppen climate zone, source water type, and sampling month were explained 25.4% (p = 0.001), 17.9% (p = 0.001), 17.8% (p = 0.001) of the total variance, respectively. Microbial communities at the POE and in the DWDS were significantly different across all utilities (PERMANOVA, p = 0.001, 999 permutations). Non-metric multidimensional scaling (NMDS) based on Bray-Curtis dissimilarity revealed clear clustering based on utilities (**Figure 1A**), independent of sampling season or source water type. Median pairwise Bray-Curtis dissimilarity for pairwise comparisons of POE and DWDS samples from different utilities was 1.0 (**Figure 1B**), indicating highly distinct community composition. Samples collected at different sites at the same utility and even at similar sampling timepoints exhibited significant divergence, indicating high level of spatial variation. For example, the utility S2 frequently showed instances of within-utility, within-month compositional divergence (Figure S1). This observation aligns with previous findings that bacterial populations in different parts of the same DWDS vary substantially which could be attributed to localized environment and regrowth dynamics ^18,52^. Of all the utilities, different sites within the utility G2 had more similar communities (average Bray-Curtis dissimilarity = 0.4) which could potentially be attributed to smaller size of the DWDS and lower water ages.

**Figure 1.**
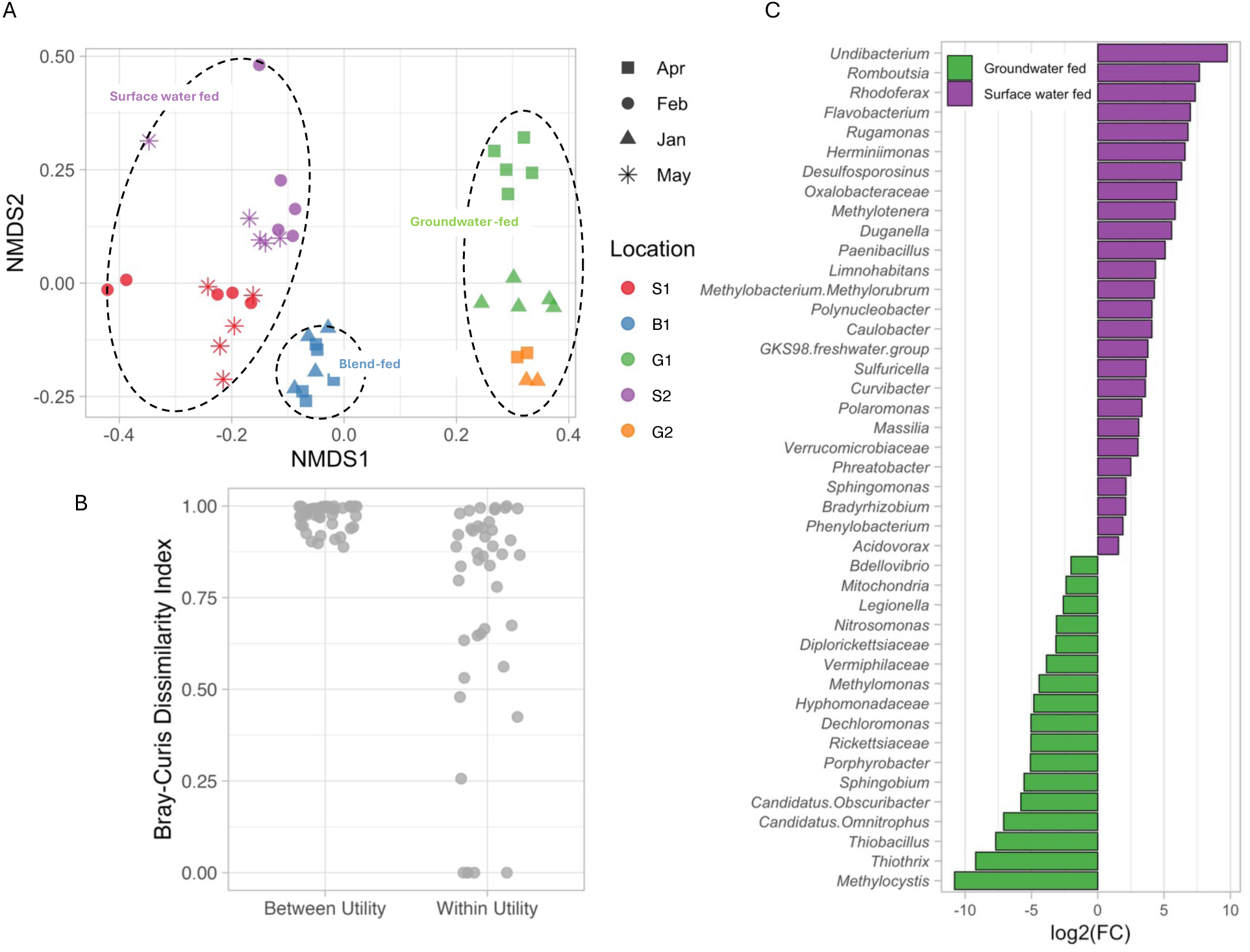
A: Non-metric multidimensional scaling (NMDS) plot of POE and DWDS samples based on Bray-Curtis dissimilarity index demonstrate utility specific as well as source water driven clustering. B: Jitter plot of pairwise Bray-Curtis dissimilarity index of lumped samples between different utilities and within the same utility, with the latter category divided into different or same sampling month, ****: T-Test p-value = 1.2e-26. C. DESeq2 analysis showing the log2fold differential abundance of the different genera between surface water and groundwater-fed systems. *: Benjamini–Hochberg adjusted p-value < 0.05. **: p-value < 0.01;*** p-value < 0.001; ****: p-value < 0.0001.

While microbial community composition was utility-specific, there was also clear clustering based on source water type for a utility (PERMANOVA, p = 0.001, 999 permutations) (**Figure 1A**) likely driven by taxa originating from groundwater or surface water environments. Of the 92 detected genera, 42 genera exhibited significant differences in abundance between surface water and groundwater as source types (**Figure 1C**). For example, *Rhodoferax*, a genus commonly found in photic aquatic environments^53^, was abundant in surface water utilities but undetectable in groundwater systems. The same pattern applied to *Romboutsia*, associated with potable water reuse^54^, and two typical surface water taxa of *Undibacterium* and *Flavobacterium* ^55,56^. In contrast, *Thiothrix*, *Methylocystis*, *and Thiobacillus* were exclusively detected in groundwater utilities. These genera have been linked to sulfur and methane-based chemolithotrophy in groundwater ecosystems ^57–59^. *Candidatus Omnitrophus*, a proposed biomarker for saline groundwater ^60^, was also found exclusively in groundwater utilities which were coastal in nature, potentially reflecting seawater influence on groundwater. Previous studies have shown that the source water influences the biological and chemical quality of drinking water in downstream locations ^61^. Multiple studies have reported that shifts in source water composition can impact microbial community composition in both planktonic^62,63^ and biofilm phases^63,64^. Notably, Learbuch et al. demonstrated that source water had greater explanatory power in terms of DWDS biofilm compositions than pipe materials, a factor commonly cited as a major determinant of biofilm composition. In this study we find that the some of the key genera differentially abundant in systems supplied by surface and ground water could be attributed to source water type.

### 3.2 Microbial cell concentrations were correlated with utility-specific water quality characteristics and DBPs concentrations

Total cell counts (TCC) and intact cell counts (ICC) across all chloraminated water samples ranged from 2.5 × 10^4^ to 3.6 × 10^5^ cells/L and 6.7 × 10^1^ to 6.1 × 10^3^ cells/L, respectively (Table S3). These values correspond to typical ranges in reported drinking water ^40,65,66^. There was a two- to three-log reduction in both TCC and ICC from source water to finished water. Seasonal variations were evident, with higher microbial cell counts detected in spring compared to winter (**Figure 2**). For instance, in utility S1, the TCC were elevated in May and showed significant positive correlations with DOC (Spearman r = 0.68, p = 0.03) and water temperature (Spearman r = 0.93, p = 0.003) (Table S4), which is consistent with previous findings that higher organic substrate availability and warmer conditions can result in microbial growth ^8,67–69^. However, this trend was not present at other utilities. For example, at utility S1, the TCC was not significantly correlated with either temperature (Spearman r = 0.54, p = 0.11) or DOC (Spearman r = 0.35, p = 0.32), while exhibiting significant correlations with sulfate (Spearman r = 0.93, p = 0.00011) and nitrate (Spearman r = -0.94, p = 0.00007). The ICC/TCC ratio was used as a proxy to examine the viable proportion of the microbial community. As expected, the DWDS had substantially lower ICC/TCC ratio (9.7 ± 5.6%) as compared to source water (67.8 ± 23.1%), indicating that the majority of cells measured as TCC were likely had damaged cell membranes. Notably, though drinking water samples collected in winter months had lower TCCs but with consistently higher ICC/TCC ratios (**Figure 2**). For example, in the utility S1, the ICC/TCC ratios declined from 9.8 ± 7.5% in February to 1.7 ± 1.7% in May despite no significant change in disinfectant residual concentrations. Total chorine residual was not significantly correlated with the cell counts in any utilities in this study. One plausible explanation is that chlorine residuals within each utility were relatively stable seasonally, while changes in TCC and ICC could be governed by changes in source water quality characteristics.

**Figure 2:**
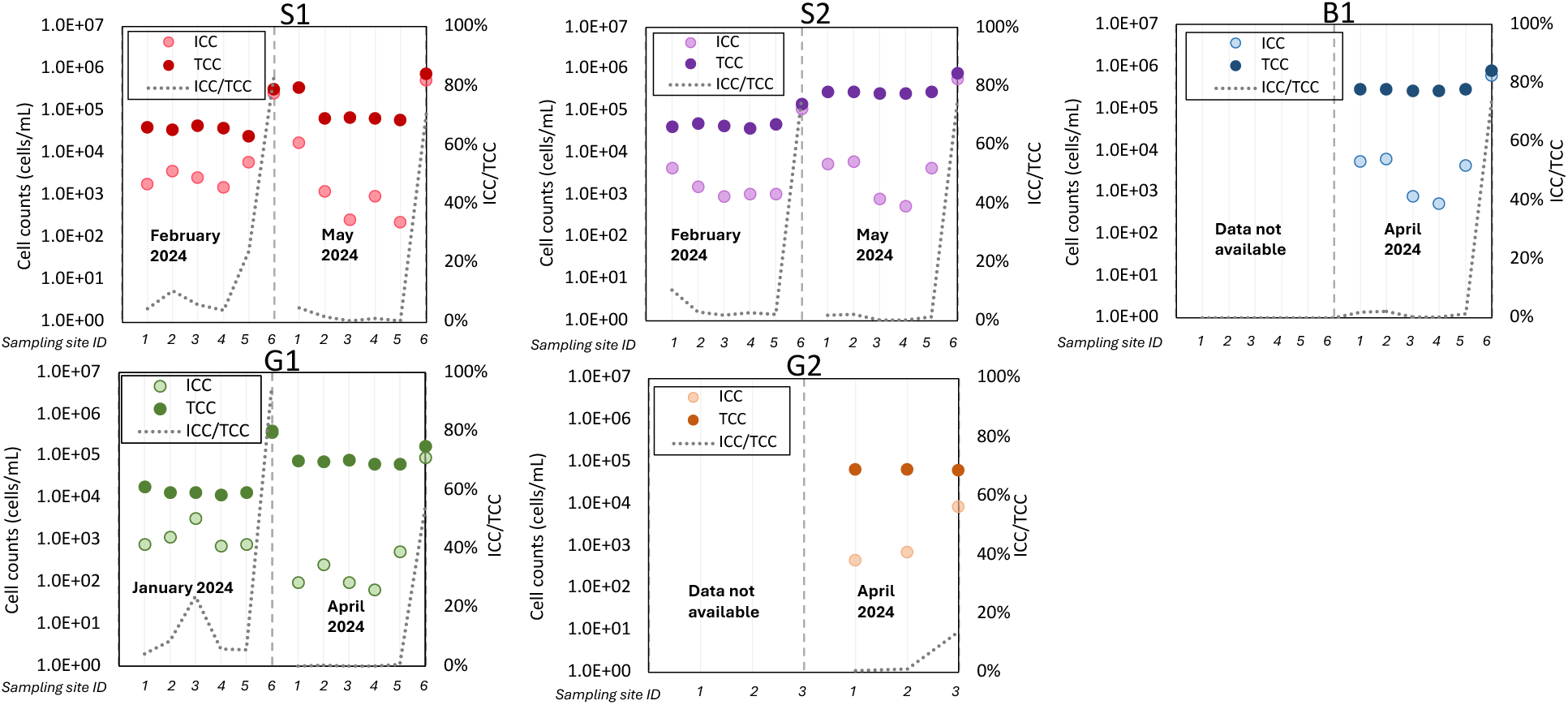
TCC, ICC, and ICC/TCC ratio per sampling site for each utility. For G2, site 1 is location in DWDS, site 2 is POE, and site 3 is source water. For all other utilities, site 1-4 are locations in DWDSs, site 5 is POE, and site 6 is source water.

A particularly noteworthy pattern emerged in the correlation between TCC and DBPs. Across all utilities, DBPs frequently exhibited strong positive correlations with TCC. For instance, THMs, the most dominant DBPs in drinking water systems ^9,70^, were consistently and positively associated with TCC (Spearman r > 0.76, p < 0.05). In the utility S1, TCCs were positively correlated with DOC (Spearman r = 0.68, p = 0.01) and temperature (Spearman r = 0.93, p = 0.01), and a contemporary study showed these factors also strongly influence DBP levels ^39^. DOC may serve a dual role, as a carbon source supporting microbial growth^71^ and as a precursor pool for DBP formation ^72^. DOC consumption was likely associated with microbial growth as reflected in the negative correlation between DOC and ICC in S1. However, this observation may not generalize to other utilities. For example, in the utility G1, TCCs increased in May despite stable water temperatures across sampling periods and this change was also uncorrelated with DOC. In utility S2, strong positive correlations were observed between TCCs and THMs and HAMs, but these trends did not correlate with either temperature or DOC. These inconsistencies raise the possibility that microbial biomass itself may contribute as DBP precursors, particularly through cellular decay or the release of microbial-derived organic matter, which can react with disinfectants to form DBPs. This hypothesis is supported by prior work showing that extracellular polymeric substances (EPS) and microbial cell lysis products can act as nitrogenous and carbonaceous DBP precursors ^73–75^. Additional research is warranted to quantify the contribution of microbial biomass and decay to DBP formation under utility-specific treatment configuration and distribution system conditions.

### 3.3 *Legionella* and *Mycobacterium* abundance were associated with microbial community composition and source water types

We examined the cell counts, Shannon diversity, and ASV distributions across source water, POE, and DWDS samples for *Legionella* and *Mycobacterium* populations (**Figure 3**). *Legionella* and *Mycobacterium* cell counts were estimated by multiplying the relative abundance of ASVs with the TCC obtained using flow cytometry. We acknowledge potential biases introduced by this approach as it assumes uniform rRNA gene copy numbers and equal DNA extraction efficiency across taxa. Despite these limitations, this approach provides a useful approximation of absolute abundances, enabling comparison across samples and utilities. Spearman correlation revealed that *Legionella* and *Mycobacterium* cell counts had weak and inconsistent correlations with individual water chemistry variables and DBP concentrations (Table S5). Nevertheless, the abundance of both genera were significantly and positively correlated with community structure and membership (Spearman p < 0.05). Samples in which *Legionella* or *Mycobacterium* ASVs were detected had significantly higher community richness, evenness, and Shannon diversity than those without detectable ASVs (Figure S2). This may suggest that *Legionella* and *Mycobacterium* likely benefit from broader ecological conditions that favor microbial growth, such as nutrient availability ^68,72^.

**Figure 3.**
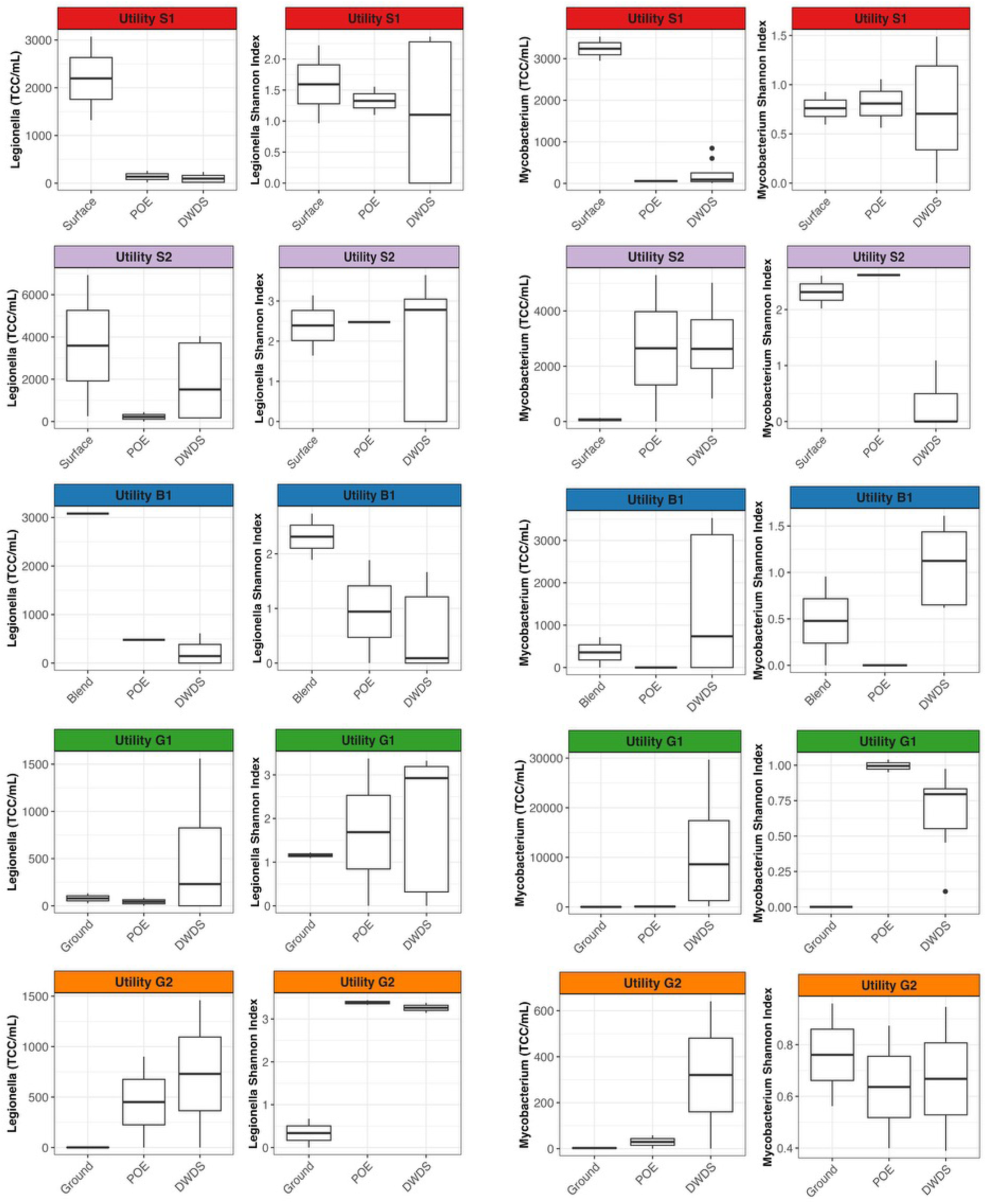
Cell concentrations (TCC/mL) and Shannon diversity of *Legionella* (left two columns) and *Mycobacterium* (right two columns) in source water, POE, and DWDS samples of all utilities.

*Legionella* cell counts decreased from source water to DWDS in surface and blend water utilities (S1, S2, and B1). In groundwater systems included in this study, *Legionella* cell abundances demonstrated an increase from source to POE and DWDS, and the Shannon diversity often peaked in DWDS samples. The ground water systems included in this study exhibited higher ambient temperatures (G1: mean DWDS temperature = 26.3 °C and G2: mean DWDS temperature = 25.9 °C). These temperatures fall within the optimal range known to support *Legionella* growth^76,77^. Also, a substantial proportion of ASVs detected in the DWDS were not detected in the upstream samples. For utilities S1, B1, and G1, *Legionella* ASVs exclusive to DWDSs accounted 67%, 73%, 62% of the total *Legionella* abundance in DWDS samples possible indicating the impact of biofilm-based seeding on *Legionella* populations in the DWDS. Unlike *Legionella*, *Mycobacterium*’s cell abundance and Shannon diversity increased in DWDSs compared to source water and POEs irrespective of source water type. The exception was the utility S1, where *Mycobacterium* concentrations in DWDS remained low. Detection and typically higher abundances of *Mycobacterium* in chloraminated drinking water systems has been previously attributed to enhanced resistance to chloramine ^78–80^. Previous studies have shown that chloramine effectively suppresses *Legionella* proliferation but is considerably less effective with respect to *Mycobacterium* ^11,19^. In this study, all utilities used chloramine, with higher total residuals observed in groundwater utilities (3.6±0.4 mg/L) compared to blend water (2.9±0.4 mg/L) or surface water utilities (1.6±0.7 mg/L). Given *Legionella*’s reported sensitivity to chloramine, the higher residual in groundwater utilities could further suppress its population abundance and diversity. For *Mycobacterium*, it is plausible that the influence of source water is masked by higher disinfectant tolerance.

Our analysis revealed that DWDSs harbor unique *Legionella* and *Mycobacterium* populations. Although source-tracking results via FEAST, consistent with previous studies, indicate that the whole microbial community in DWDSs is largely shaped by POE water ^81^, this trend did not extend to *Legionella* and *Mycobacterium* populations. A substantial proportion of *Legionella* and *Mycobacterium* ASVs were found exclusive in DWDSs, with little to no overlap with upstream POE and/or source waters. Interestingly, this community divergence was not associated with water age. For example, in utility B1, with a lower average water age (44 hours), had 100% unique *Mycobacterium* ASVs in DWDS, compared to only 28% in utility S1, where the average water age was longer (72 hours). Given that microbial taxa in DWDS ultimately originate from upstream, this divergence in trends between overall community and *Legionella* and *Mycobacterium* populations may reflect preferential growth (and subsequent detachment) of the latter in biofilms. Indeed, the differential abundance analysis indicated that several genera linked to drinking water biofilms were significantly more enriched in DWDS than POE samples, including *Aquabacterium* ^82,83^, *Phreatobacter* ^84^, as well as *Gallionella* commonly found in cast iron pipe biofilms ^85,86^ (Table S6).

### 3.4 *Legionella* and *Mycobacterium* ASV’s show marker geographic structuring

We examined the spatial and temporal structuring of *Legionella* and *Mycobacterium* populations using pairwise Bray-Curtis dissimilarity (Figure S3). Overall, dissimilarity declined with progressively finer spatial and temporal scales, indicating that *Legionella* and *Mycobacterium* populations become more similar when samples were collected in closer proximity in time and location. Comparisons between different utilities showed dissimilarity saturation (Bray-Curtis index of 1.0), reflecting that *Legionella* and *Mycobacterium* populations from separate utilities share few to no ASVs. This lack of overlap highlights strong geographic structuring, consistent with the broader trends observed for the entire microbial community (**Figure 1**). Principal-coordinate analysis (PCoA) of *Legionella* and *Mycobacterium* ASVs revealed a horseshoe effect (Figure S4) ^88^, a common artifact that arises when distance metrics that saturate^89^. This pattern reinforces that *Legionella* and *Mycobacterium* populations diverge significantly across utilities with limited overlap.

To further explore the community divergence among utilities, we selected the top 10 abundant *Legionella* and *Mycobacterium* ASVs from each utility. These top ASVs accounted for a substantial proportion of the overall population: 68–79% of total *Legionella* abundance and 81–98% of total *Mycobacterium* abundance. Many of the dominant ASVs observed in DWDSs were undetected in the source water, likely due to their low abundance. Nevertheless, the influence of source water is reflected by the PCoA analysis, which showed clear separation by source water type (PERMANOVA, *p* < 0.01) (Figure S4). Further, PCoA-based on UniFrac distance, which accounts for phylogenetic relatedness, revealed distinct clustering patterns based on source water types: groundwater-fed utility samples tended to cluster tightly, while surface water-fed samples were more dispersed (Figures S5 and S6). This indicates that groundwater systems may harbor more phylogenetically related *Legionella* and *Mycobacterium* strains compared to surface water systems. Even when similar ASVs are present in multiple raw water sources, differences in treatment processes and DWDS environments may drive population divergence. For instance, dominant *Legionella* and *Mycobacterium* ASVs were utility-specific (**Figure 4** and **Figure 5**). For example, *Mycobacterium iranicum-like* ASV281 was the most abundant ASV in utility G1, with the single ASV accounted for 32.7 % of the total *Mycobacterium* abundance, but it was absent from all other utilities. Similarly, 42 out of 45 of the most abundant *Legionella* ASVs appeared to be restricted to single utilities. Although *Mycobacterium* and *Legionella* are frequently detected in municipal drinking water systems, these results reveal that their compositions is distinct across utilities. These patterns point to strong utility-specific selective pressures shaping microdiversity, with important implications for designing targeted microbial control strategies and supporting utility-specific water safety management. Several ASVs were also identified in multiple utilities. For example*, Legionella* ASV993 was found in both S1 and S2, and *Mycobacterium mucogenicum*-like ASV44 appeared in B1, G1, and G2. However, these cases were rare: only 3 of the top 45 *Legionella* ASVs were detected in more than one utility. *Mycobacterium* showed slightly broader distribution, with 6 of the top 14 ASVs shared across utilities. Identifying the key environmental and operation-related variables that drive these patterns will be critical for understanding the mechanisms underlying strain-level divergence in drinking water systems.

**Figure 4.**
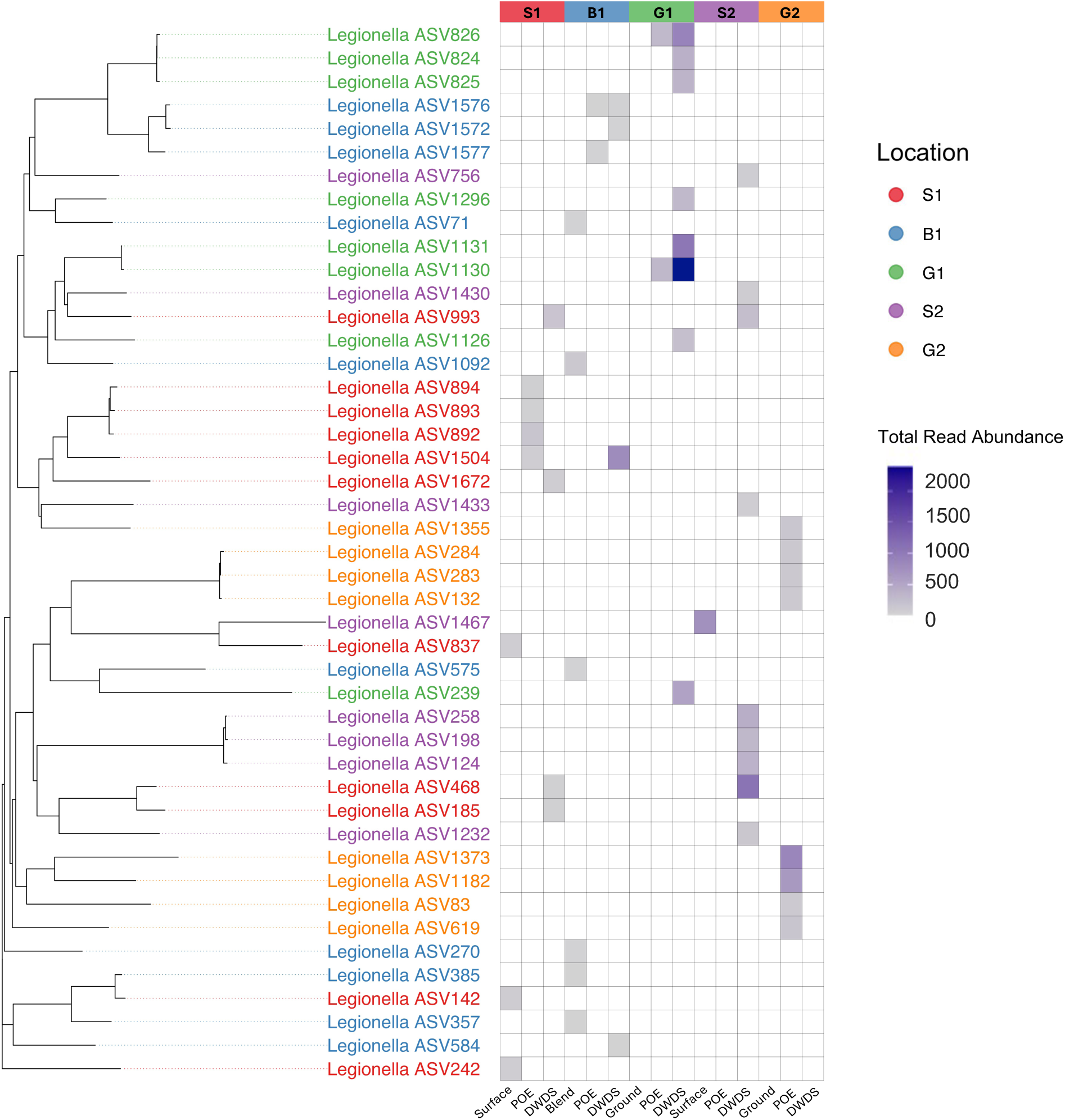
Phylogenetic tree of the top 10 *Legionella* ASVs from each utility. Each ASVs were colored based on the utility. The heatmap shows ASV read abundances across source water, point of entry (POE), and distribution system samples (DWDS).

**Figure 5.**
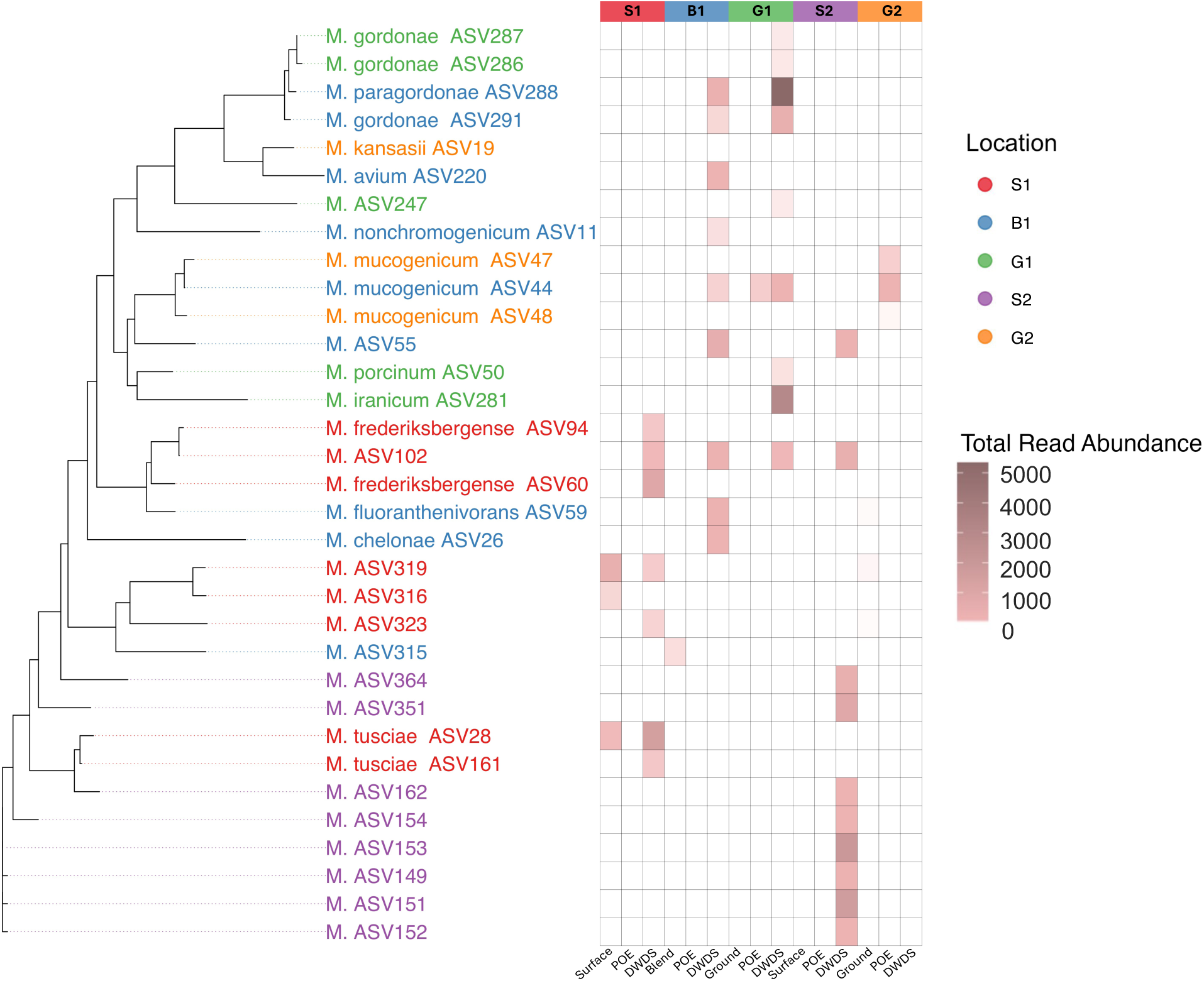
Phylogenetic tree of the top 10 Mycobacterium ASVs from each utility. Each ASVs were colored based on the utility. The heatmap shows ASV read abundances across source water, point of entry (POE), and distribution system samples (DWDS).

## Acknowledgments

This research was supported by USEPA Grant R840606.

## Data Availability

The raw 16S rRNA gene sequences are available on NCBI at BioProject accession number PRJNA1393449. All codes for data analysis and visualization are available upon request.

## Supporting Information

**Figure S1:**
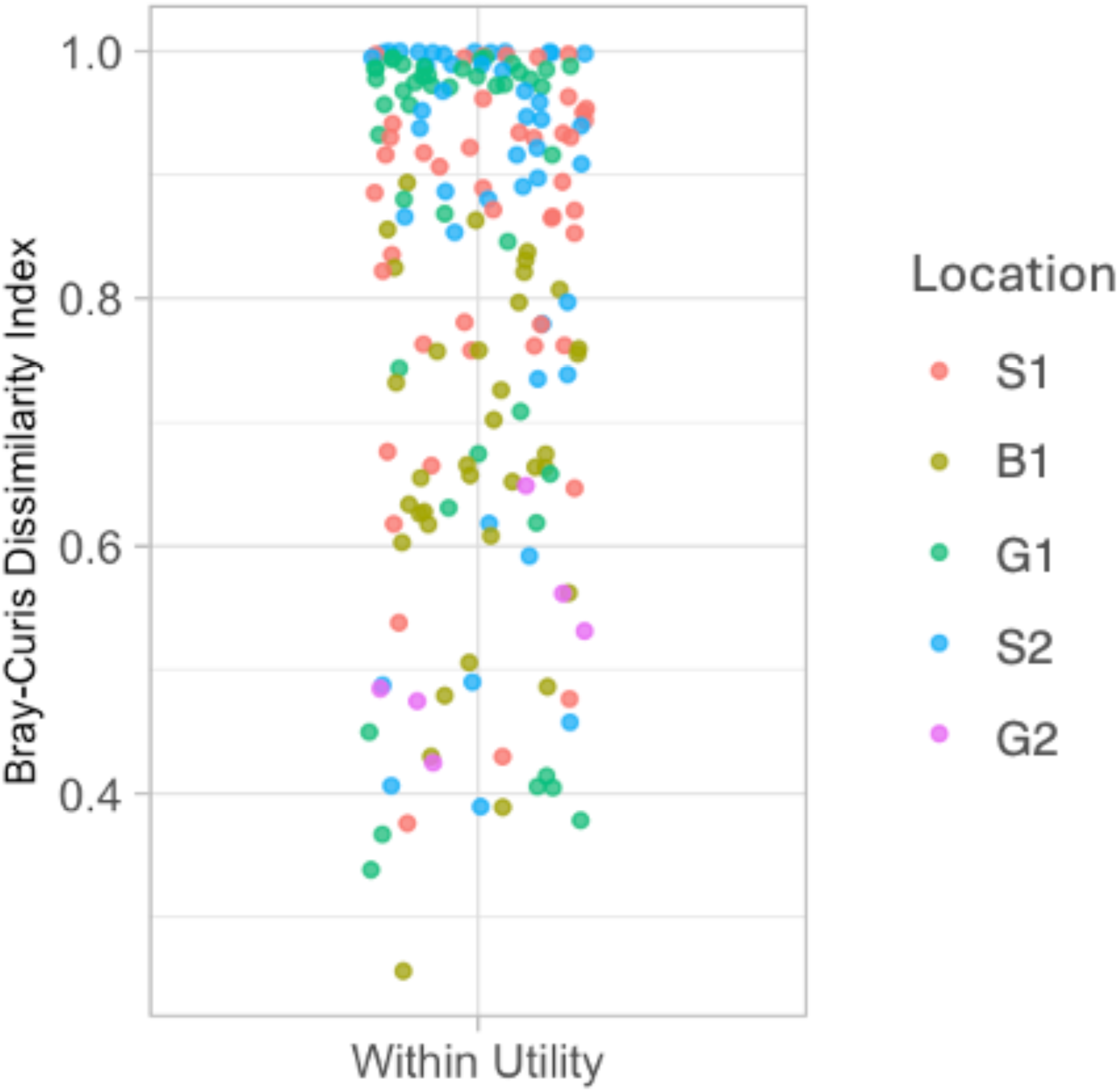
Bray-Curtis dissimilarity index for pairwise comparisons of POE and DWDS samples

**Figure S2.**
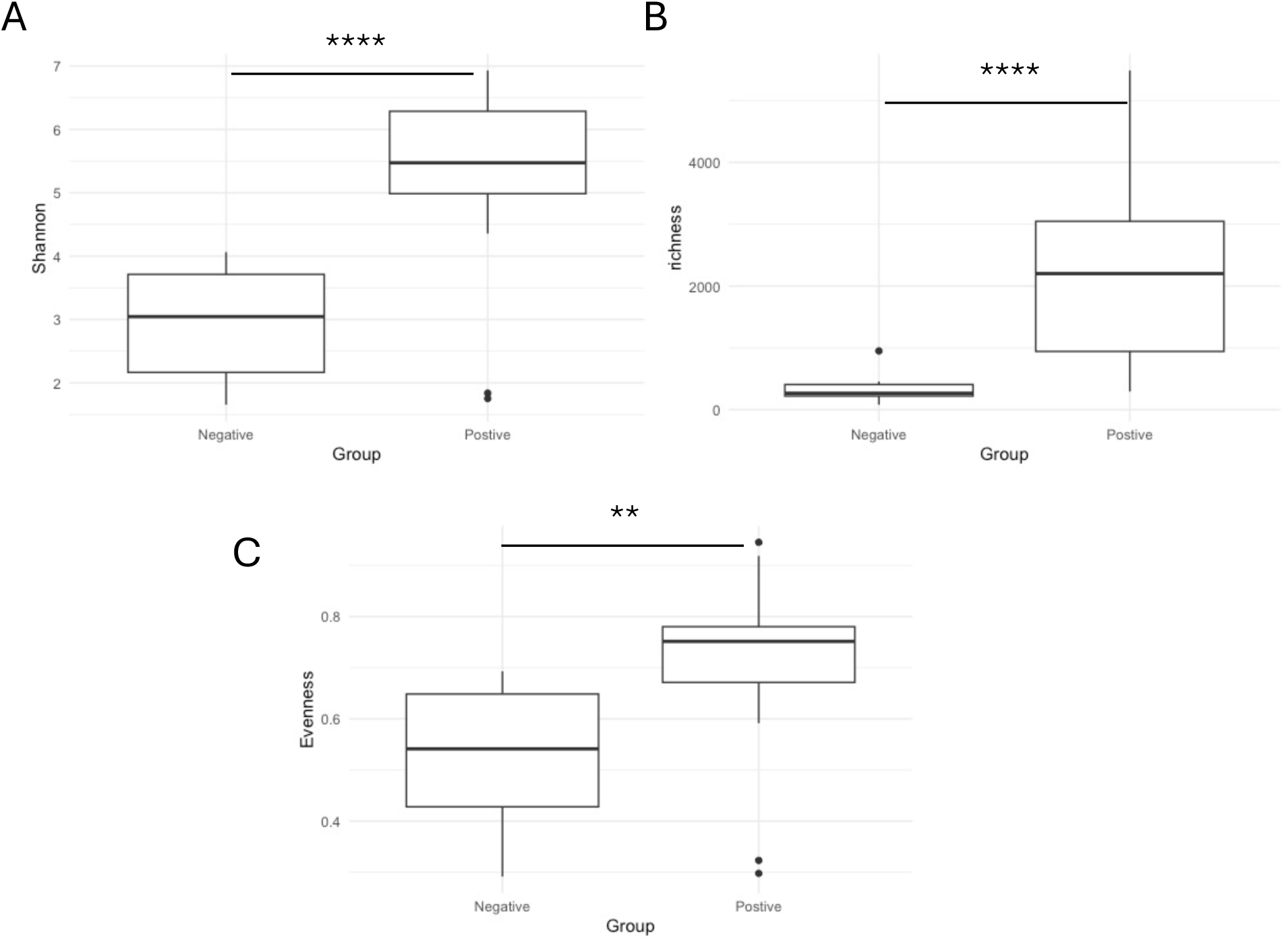
Microbial Community’s Shannon diversity (A), richness (B), and Evenness (C) in samples with detected *Legionella* or *Mycobacterium* ASVs (noted as “positive”) and without detected ASVs (noted as “negative”). **: T. test p-value < 0.01;*** p-value < 0.001; ****: p-value < 0.0001.

**Figure S3.**
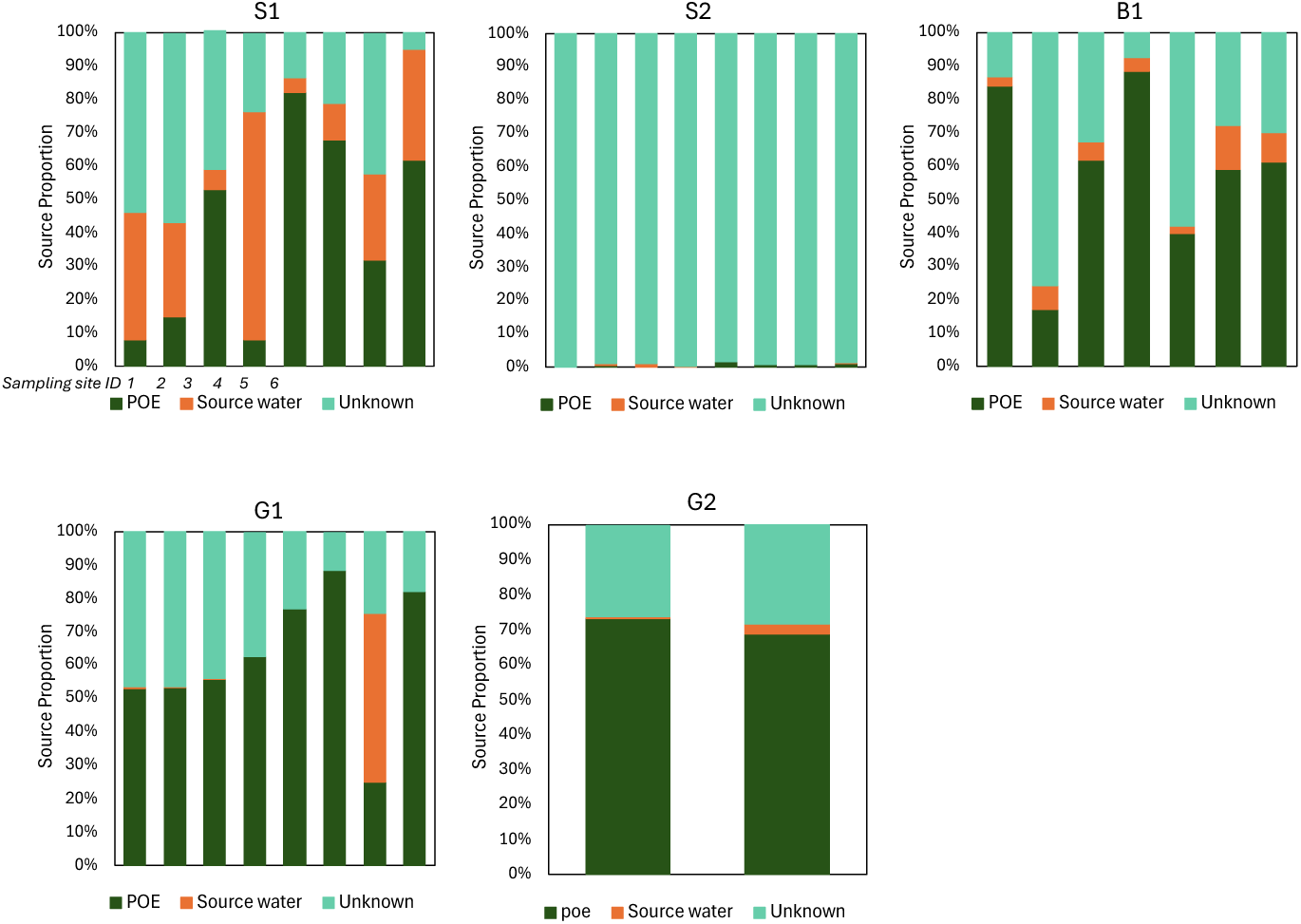
Source estimates considering point-of-entry and raw source water for drinking water distribution systems.

**Figure S4.**
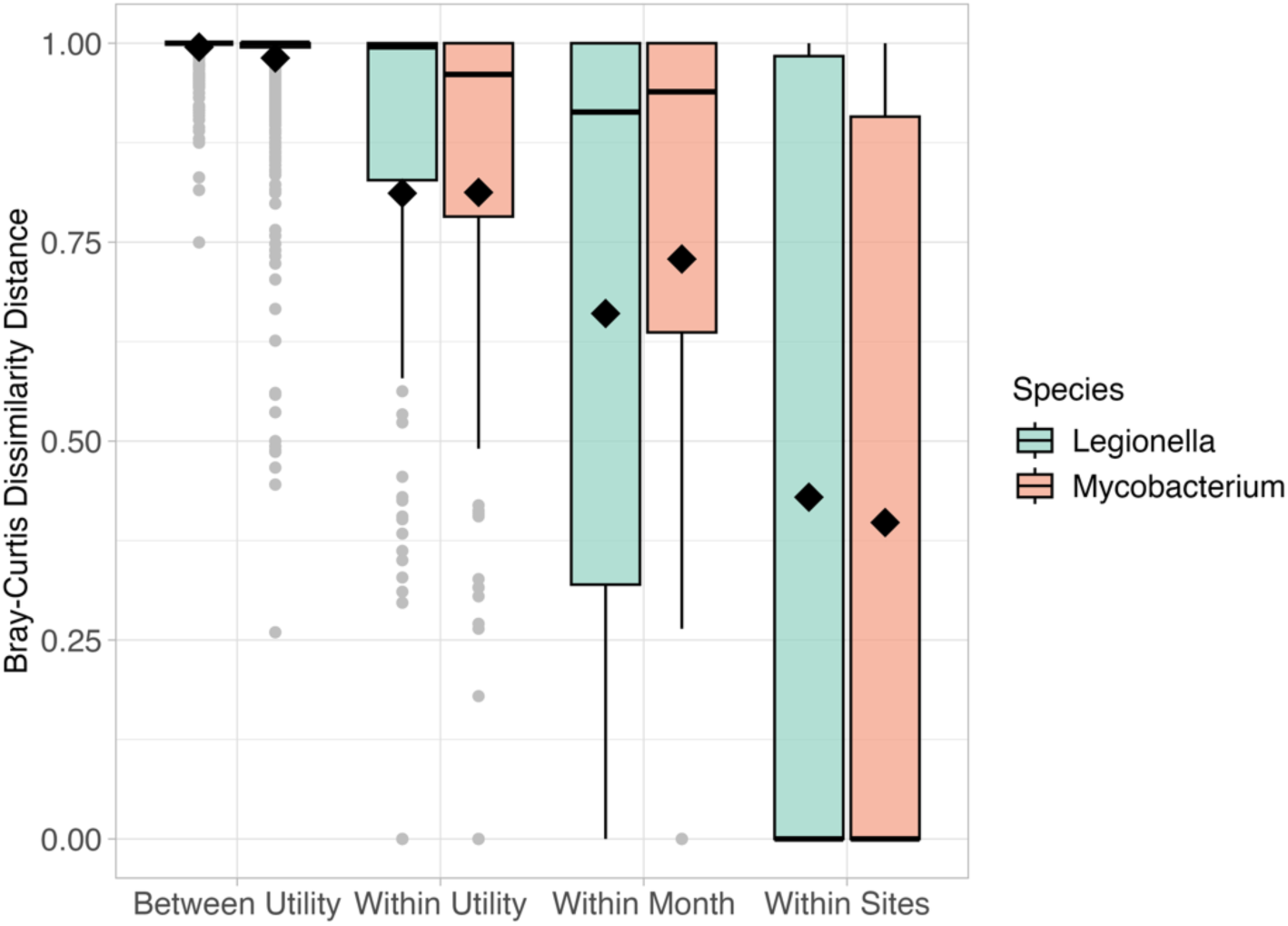
Pairwise Bray-Curtis dissimilarity index of Legionella and Mycobacterium calculated for samples grouped across progressively finer spatial and temporal scales.

**Figure S5.**
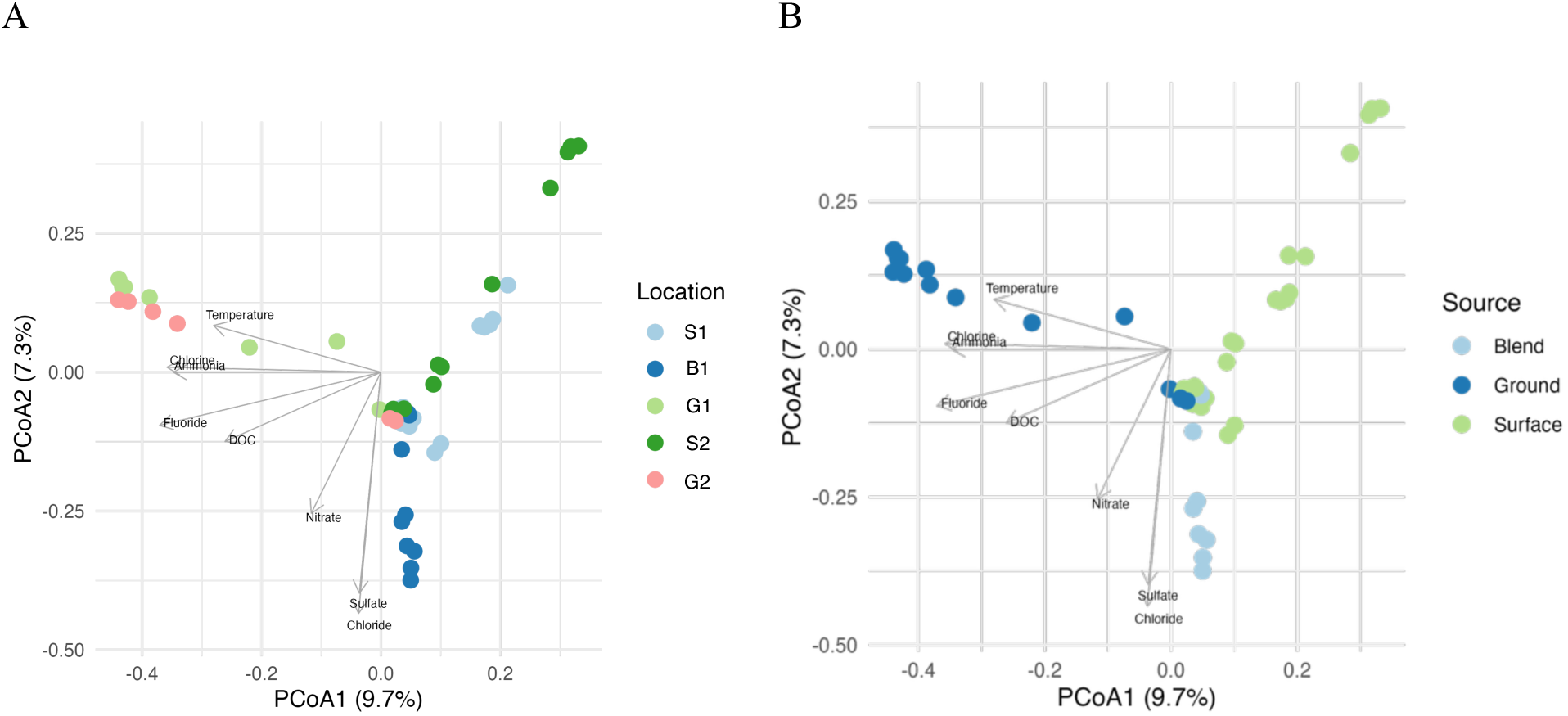
Principal-coordinate analysis (PCoA) of *Legionella* and *Mycobacterium* ASVs based on A: different locations (i.e., utility IDs) and B: source water types.

**Figure S6.**
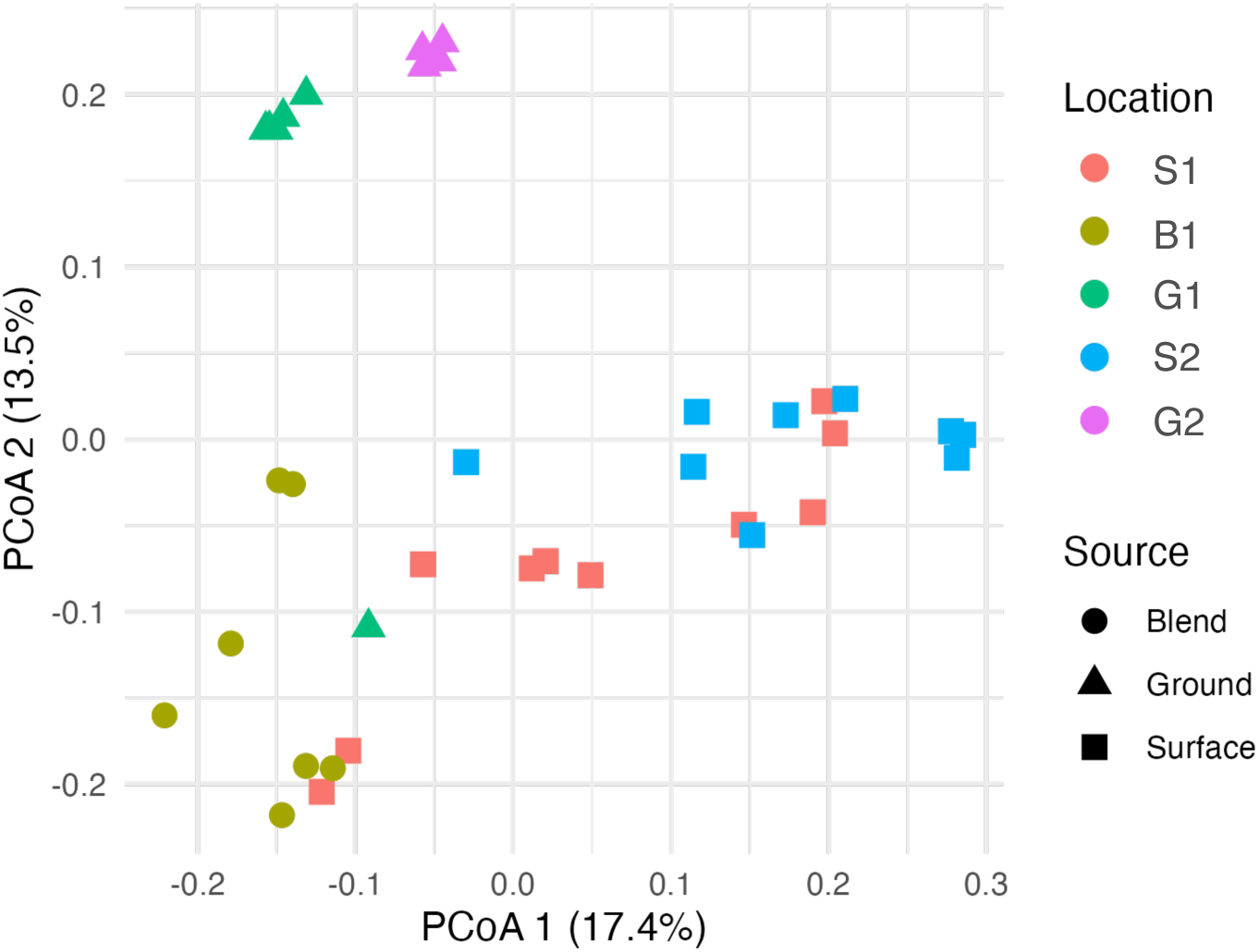
Principal-coordinate analysis (PCoA) with distance metrics of Unifrac (weighted) for *Legionella* ASVs based on different locations (i.e., utility IDs) and source water types.

**Figure S7.**
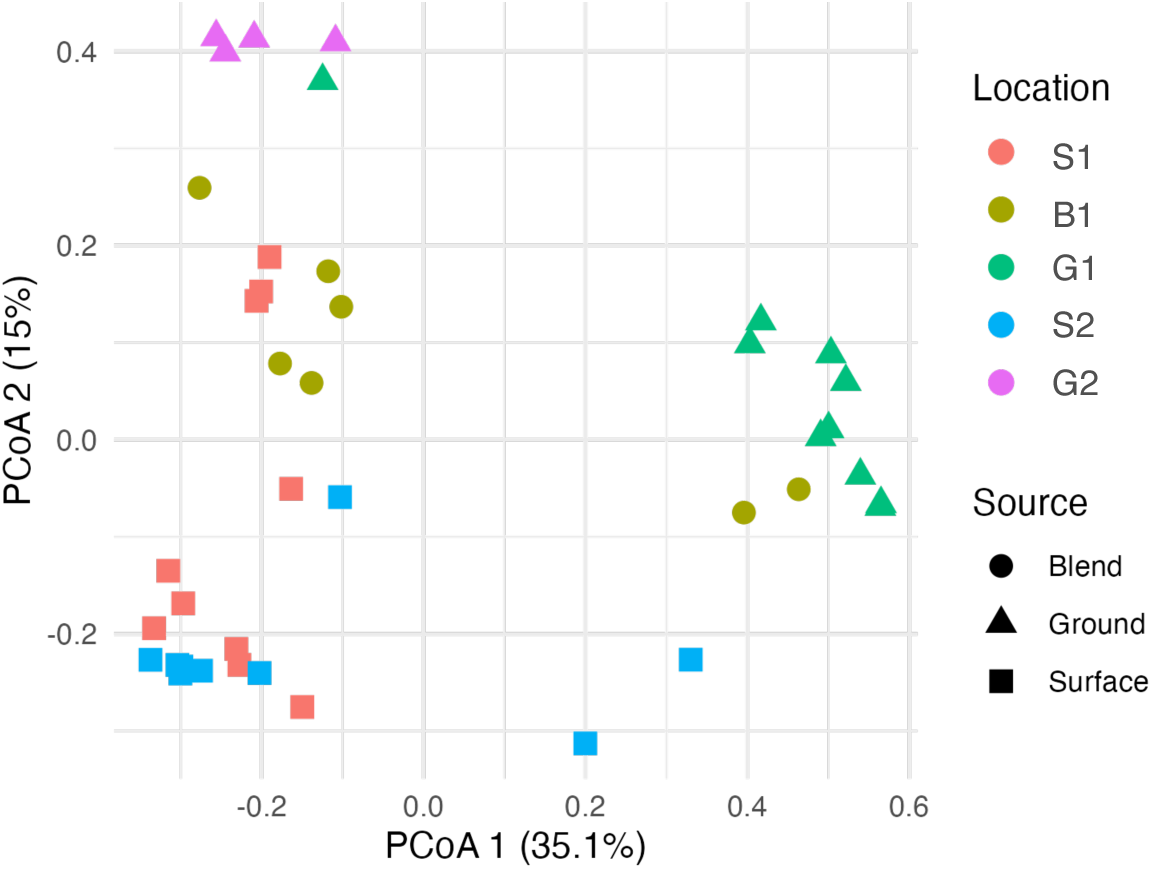
Principal-coordinate analysis (PCoA) with distance metrics of Unifrac (weighted) for *Mycobacterium* ASVs based on different locations (i.e., utility IDs) and source water types.

